# LVentiView: An Open-Source Software for Automated 3D Left Ventricular Mesh Reconstruction and Analysis from Cardiac MRI

**DOI:** 10.64898/2026.05.22.727166

**Authors:** Ina Braun, Yong Wang, Alexander S. Ecker, Eberhard Bodenschatz

## Abstract

Patient-specific cardiac modeling requires accurate three-dimensional representations of the left ventricle (LV) reconstructed from cardiac magnetic resonance imaging (MRI). Here, we present LVentiView, an open-source software that bridges medical imaging and cardiac simulation by automating the full pipeline from MRI segmentation to simulation-ready volumetric meshes, with integrated tools for volumetric analysis and regional myocardial thickness calculation. We validate LVentiView on the Sunnybrook Cardiac Dataset, comprising healthy subjects and three cardiac pathologies. LVentiView achieves blood pool segmentation at the inter-expert level. The generated meshes are verified by comparing LV volumes extracted from the meshes to those computed from expert manual segmentation masks, with volumes and cardiac parameters agreeing within inter-expert variability across all four cardiac pathologies. In addition, mesh-derived regional thickness maps capture pathology-specific patterns, including wall thickening in hypertrophic cases. LVentiView is freely available on GitHub and provides an accessible, validated foundation for patient-specific cardiac modeling.

**Highlights:**

- LVentiView automates the full pipeline from MRI segmentation to simulation-ready meshes.
- Mesh-derived cardiac volumes and parameters match expert manual segmentation accuracy.
- Thickness maps capture pathology-specific patterns, validating geometrical fidelity.
- Segmentation runs at ≈ 0.07 s per slice; meshing under 3 min per frame.

**Graphical Abstract:** 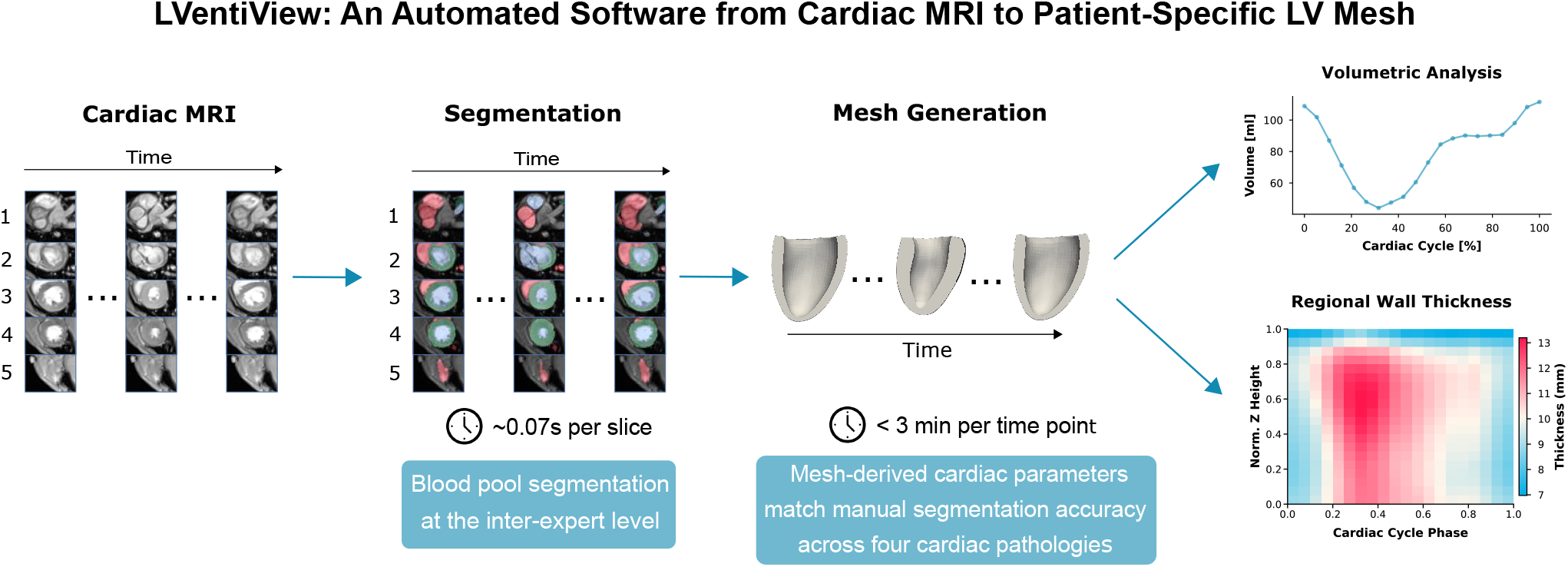

## 1. Introduction

Before cardiac imaging became routine in clinics, cardiac function was primarily assessed through pressure and flow measurements. With the advent of advanced imaging techniques in the early 1960s, such as left ventriculography, clinicians began using left ventricular cavity volume (LVV) and ejection fraction (EF), the latter is defined as the stroke volume divided by the end-diastolic volume (Folse and Braunwald, 1962; Kennedy, Baxley, Figley, Dodge and Blackmon, 1966). Today, EF is a cornerstone of cardiology routinely applied in diagnosis and prognosis (Marwick, 2018).

While global volumetric indices such as LVV and EF are the clinical standard, they do not capture the full threedimensional geometry of the left ventricle, and therefore do not allow assessment of local parameters such as regional myocardial wall thickness. Such local parameters are clinically relevant, as wall thickness serves as a diagnostic marker for left ventricular hypertrophy (Baltabaeva, Marciniak, Bijnens, Moggridge, He, Antonios, MacGregor and Sutherland, 2008), contributes to diagnosing heart failure with preserved EF (Pieske, Tschöpe, De Boer, Fraser, Anker, Donal, Edelmann, Fu, Guazzi, Lam et al., 2019), and assists in detecting infarcted regions (Lieberman, Weiss, Jugdutt, Becker, Bulkley, Garrison, Hutchins, Kallman and Weisfeldt, 1981). Beyond clinical diagnosis, three-dimensional models of the left ventricle (LV) are becoming increasingly important for patient-specific biomechanical simulations of cardiac mechanics (Dabiri, Van der Velden, Sack, Choy, Guccione and Kassab, 2020; Hadjicharalambous, Stoeck, Weisskopf, Cesarovic, Ioannou, Vavourakis and Nordsletten, 2021; Miller, Kerfoot, Mauger, Ismail, Young and Nordsletten, 2021; Mu, Chan and Yap, 2025), further motivating the need for accurate and accessible 3D LV reconstruction from cardiac magnetic resonance imaging (MRI).

Existing tools address part of the workflow necessary to reconstruct a 3D LV mesh from cardiac MRI. 3D Slicer (Fedorov, Beichel, Kalpathy-Cramer, Finet, Fillion-Robin, Pujol, Bauer, Jennings, Fennessy, Sonka et al., 2012) and ITK-SNAP (Yushkevich, Piven, Cody Hazlett, Gimpel Smith, Ho, Gee and Gerig, 2006) provide powerful tools for segmenting 3D and 4D MRI series, but do not produce simulation-ready meshes. Simulation frameworks such as lifex (Africa, 2022) allow biomechanical simulations, but they lack a pipeline to create simulation ready meshes from cardiac MRI. SimVascular (Updegrove, Wilson, Merkow, Lan, Marsden and Shadden, 2017) offers an integrated workflow from medical imaging to simulation, but targets cardiovascular fluid dynamics rather than solid-mechanics cardiac modeling. LVentiView fills the gap between medical imaging software and solid-mechanics cardiac simulations, by providing an automated, open-source workflow from MRI to volumetric mesh, complemented by integrated tools for volumetric analysis and regional myocardial thickness calculation. LVentiView is implemented for short-axis (SAX) MRI series, and is freely available on GitHub^1^.

The contributions of this paper can be summarized as follows:

a. We introduce LVentiView, an open-source framework that bridges medical imaging and cardiac simulation, automating the full pipeline from MRI segmentation to simulation-ready volumetric meshes, complemented by tools for volumetric analysis and regional myocardial thickness calculation.
b. The generated meshes are verified by comparing LVVs extracted from the meshes against LVVs from expert manual segmentations, with volumes and cardiac parameters agreeing within inter-expert variability across all four cardiac pathologies.
c. Regional myocardial thickness maps capture pathology-specific remodeling patterns, including wall thickening in hypertrophic cases.

## 2. Related Work

### 2.1. Patient-Specific Cardiac Simulations

Biomechanical simulations of cardiac mechanics increasingly shift from idealized to patient-specific left ventricular geometries, both in finite-element method (FEM) simulations (Guccione, Costa and McCulloch, 1995; Hadjicharalambous et al., 2021; Miller et al., 2021) and in machine learning approaches, where networks are trained either on finite-element simulation data (Dabiri et al., 2020) or using physics-informed frameworks that embed the governing equations of cardiac mechanics directly into the learning process (Mu et al., 2025). For both patient-specific FEM based and neural network based simulations, generating accurate patient-specific volumetric meshes from clinical MRI data is a prerequisite.

### 2.2. Mesh Generation

To generate three-dimensional LV representations, techniques have been developed that map a continuous 3D model to segmented LV contours, producing either surface (Avendi, Kheradvar and Jafarkhani, 2016; Villard, Grau and Zacur, 2018) or volumetric meshes (Joyce, Buoso, Stoeck and Kozerke, 2022). In particular, Joyce et al. (2022) proposed a method in which a volumetric mesh is fit to segmentation masks derived from SAX MRI series. This method forms the basis of the Mesh Generation Module in LVentiView.

In this approach, the volumetric mesh is expressed as a linear combination of *n* modes, each a vector of length 3 ⋅ *n*_nodes_, where *n*_nodes_ denotes the number of mesh nodes. These modes were derived via Proper Orthogonal Decomposition (POD) from manually reconstructed 3D LV geometries derived from MRI data from the Multi-Modality Whole Heart Segmentation (MM-WHS) dataset Zhuang and Shen (2016a,b). The dataset includes both healthy subjects and subjects diagnosed with diseases such as heart failure with infarction and dilated cardiomyopathy Zhuang and Shen (2016a,b). The POD basis captures size and shape variation independently. This means that scaling a heart to a different size corresponds exactly to multiplying its mode coefficients by the same scaling factor.

The mesh is fit to *k* spatially aligned 2D MRI slices,with corrective shifts applied in the SAX plane to account for patient motion. Fitting is performed by optimizing four sets of parameters: mode coefficients *α*, global mesh position *γ*, global mesh rotation *δ*, and per-slice corrective shifts *β*. The parameters are estimated using the Adam optimizer (Kingma, 2014), minimizing a loss function that combines Dice-based overlap between the myocardial region (*Y*^myp^ ) and blood pool region (*Y*^bp^ ) of the sliced mesh, and the corresponding segmentation masks (*X*^myo^, *X* ^bp^). The loss further includes regularization *l*_1_ (*θ*) = ∑ _*j*_ |*θ*_*j*_ | = to penalize large parameter values. The resulting loss function is given by

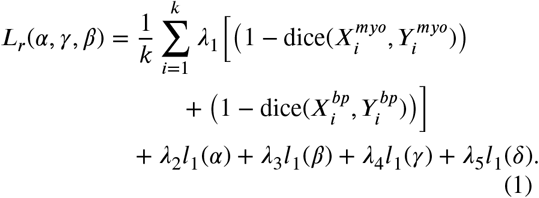

The Dice coefficient between segmentation masks *A* and the sliced mesh *B* is defined as

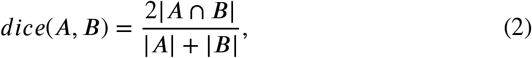

where |*X* |denotes the area of contour *X*.

### 2.3. Segmentation

Mesh fitting requires segmentation masks as input. Manual segmentation of the LV boundaries is prone to interobserver variability. Even among highly experienced annotators, the average Dice overlap, which quantifies the agreement between two segmentation masks, is 93% for the LV blood cavity and 88% for the myocardium (Bai, Sinclair, Tarroni, Oktay, Rajchl, Vaillant, Lee, Aung, Lukaschuk, Sanghvi et al., 2018). Automated methods based on convolutional neural networks (CNNs) have emerged as a reliable alternative to manual segmentation (Bartoli, Fournel, Bentatou, Habib, Lalande, Bernard, Boussel, Pontana, Dacher, Ghattas et al., 2020; Joyce et al., 2022; Koo, Lee, Ko, Lee, Kang, Kim and Yang, 2020; Wang, Zhang, Wen, Tian, Kao, Liu, Xuan, Ordovas, Saloner and Liu, 2021). These approaches produce results comparable to expert manual segmentation, with reported Dice similarity coefficients for the LV blood cavity up to 96% (Bartoli et al., 2020) and Dice similarity coefficients for the LV myocardium segmentation around 88%–89% (Bartoli et al., 2020; Koo et al., 2020).

In particular, Joyce et al. (2022) published a pretrained segmentation model for SAX slices. The model is based on the Ternaus-UNet architecture originally described by Iglovikov and Shvets (2018), which modifies the standard U-Net (Ronneberger, Fischer and Brox, 2015) with a VGG11 encoder (Simonyan and Zisserman, 2014). Joyce et al. (2022) trained the network on cardiac MRI datasets [Automated Cardiac Diagnosis Challenge (ACDC): Bernard, Lalande, Zotti et al. (2017); Bernard, Lalande, Zotti, Cervenansky, Yang, Heng, Cetin, Lekadir, Camara, Ballester et al. (2018) and Multi-Modality Whole Heart Segmentation (MM-WHS): Zhuang and Shen (2016a,b)] and fine-tuned them using a semi-supervised approach, where predicted segmentation on additional MRI data from the ACDC and Kaggle dataset (Kaggle, 2016) were used as pseudo-ground truth. These datasets include both healthy subjects and patients with a range of cardiac diseases, such as heart failure with infarction, dilated cardiomyopathy (heart failure without infarction), and hypertrophy. The Segmentation Module of LVentiView uses these pretrained models to classify each pixel in a cardiac MRI image into one of four classes: background, left ventricular myocardium, left ventricular blood pool, and right ventricle. The segmentation performance of this module as integrated into LVentiView is quantified in Section 4.1.1.

Previous studies have developed tools addressing individual steps in going from medical images to simulation-ready 3D representations: segmentation platforms such as 3D Slicer and ITK-SNAP (Fedorov et al., 2012; Yushkevich et al., 2006), and surface or mesh reconstruction methods (Avendi et al., 2016; Joyce et al., 2022; Villard et al., 2018) in isolation. However, to our knowledge no available software consolidates these steps into a single, automated pipeline with a graphical user interface (GUI) that takes cardiac MRI as input and produces simulation-ready LV meshes as output. LVentiView fills this gap between medical imaging software and cardiac simulation by providing an automated, open-source framework with a graphical user-interface covering the full pipeline from MRI segmentation to volumetric mesh generation, complemented by integrated tools for volumetric analysis and regional myocardial thickness calculation.

## 3. Methodology

### 3.1. LVentiView Overview

The workflow of LVentiView consists of two modules. The Segmentation Module segments the blood pool, myocardium and right ventricle in input cardiac MRI images. The Mesh Generation Module fits a low-dimensional basis to the segmentation masks to reconstruct a 3D volumetric mesh of the LV that captures the full cardiac cycle. Additionally, the blood pool and myocardium volume as well as regional myocardial wall thickness maps are calculated from the generated meshes.

LVentiView can be run directly from the terminal or via a GUI. Upon opening the GUI, users are presented with a start page providing navigation to both the Segmentation Module and the Mesh Generation Module, as shown in Supplementary Figure S1.

### 3.2. LVentiView – Segmentation Module

The Segmentation Module of LVentiView enables the user to automatically segment SAX MRI slices using the Teranus UNet pretrained by Joyce et al. (2022).

The Segmentation Module, GUI in Supplementary Figure S2, allows users to upload and process cardiac MRI series with a single click. In the default workflow (Fig. 1), users select an input folder containing the MRI images and specify an output directory. Upon initiating segmentation via the *Run Segmentation* button, segmentation masks are generated automatically. To ensure accurate downstream LVV and EF calculation the segmented MRI images are cleaned, as described below. Both the complete and cleaned segmentation masks are saved in a serialized (.pkl) format.

**Figure 1:**
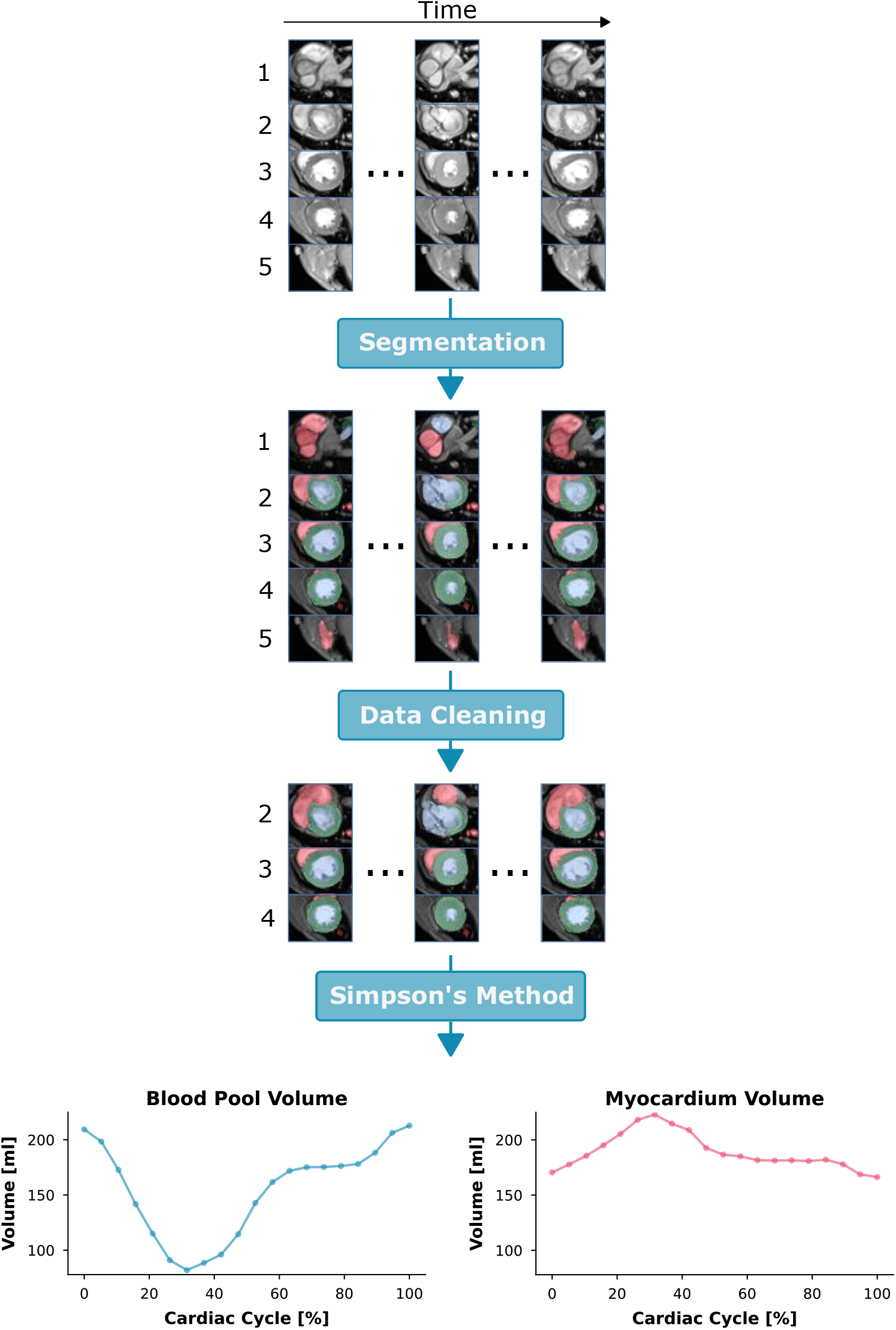
Workflow of the LVentiView Segmentation Module. **Top**: Short-axis MRI z-stack images acquired over the cardiac cycle. **Upper middle**: Segmentation masks predicted by the TeranusUNet model (blue: blood pool, green: myocardium, red: right ventricle). **Lower middle**: Cleaned segmentation stacks after removal of incomplete time frames and slices located above the mitral valve plane or below the ventricular apex. **Bottom**: Quantitative outputs of the Segmentation Module, including blood pool and myocardial volumes computed using Simpson’s method.

In the first step of data cleaning, MRI images are automatically cropped around the center of the LV. Since the segmentation networks operate on a graphics processing unit (GPU) and GPU memory is limited, the crop is chosen as the smallest square that still contains the entire LV. This is achieved by computing the average LV diameter and multiplying by a padding factor (default 2). The padding factor was manually increased when the LV extended beyond the cropped region and decreased when the complete MRI time series did not fit in the GPU memory (e.g., to 3 or 1.6).

In the next data cleaning step incomplete time points and slices outside the anatomical extent of the LV are excluded.

Specifically:

- Time points with more than two missing image planes are excluded.
- Slices above the mitral valve plane are excluded based on two morphological criteria: (i) the myocardium no longer forms a closed contour around the blood pool, and (ii) the segmentation masks of the right ventricle (RV) splits into more than three disconnected regions. A slice is removed only if both criteria agree for more than *p*_valve_ = 70% of time frames. This threshold was chosen to retain slices where the myocardium forms a closed contour during diastole but an open contour during systole.
- Apical slices are excluded if no pixels are segmented as LV myocardium or blood pool in more than *p*_apex_ = 20% of frames.

After cleaning, the myocardial and blood pool volumes are calculated from the segmentation masks using Simpson’s method, which estimates LV volume by summing the respective areas in the segmentation masks and multiplying by the inter-slice distance O’Dell (2019).

The workflow is fully automated by default, but users can adjust cleaning thresholds and enable/disable data cleaning or volume calculation.

### 3.3. LVentiView – Mesh Generation Module

To provide a full 3D geometric representation of the left ventricle for cardiac simulations and local diagnostic markers such as regional myocardial wall thickness, a lowdimensional basis is fitted to the segmentation masks (Joyce et al., 2022). This mesh fitting algorithm is implemented in the Mesh Generation Module of LVentiView (Supplementary Fig. S3). Additionally, the Mesh Generation Module enables calculation of the local myocardial wall thickness from the generated meshes.

To fit the low-dimensional basis to the segmentation masks the loss function in Eq. 1 is minimized. The weighting coefficients in the loss function (Eq. 1) were determined via hyperparameter search as *λ*_1_ = 5, *λ*_2_ = 10^−9^, *λ*_3_ = 10, *λ*_4_ = 0.3, and *λ*_5_ = 1.

During mesh fitting, optimization is performed independently for each time point for 10,000 iterations, and the mesh achieving the highest average Dice score across both blood pool and myocardium over all fitting iterations is selected, yielding an average myocardial Dice of 0.84 ± 0.05 and an average blood pool Dice of 0.91 ± 0.04. All weighting coefficients and the maximum number of fitting iterations can be adjusted by the user in the GUI. In the default workflow (Fig. 2), users select a folder containing segmented MRI data in a serialized (.pkl) format and an output directory for the meshes and analysis results. By clicking the *Run Mesh Fitting* button, the software constructs one volumetric mesh per imaged time step, saves the meshes in .vtk format, and generates visualizations of the mesh overlaid on the MRI images along with Dice scores quantifying the fit to the segmentation masks. To speed up mesh generation, mesh fitting is performed independently for each time point and parallelized across time points on the GPU.

**Figure 2:**
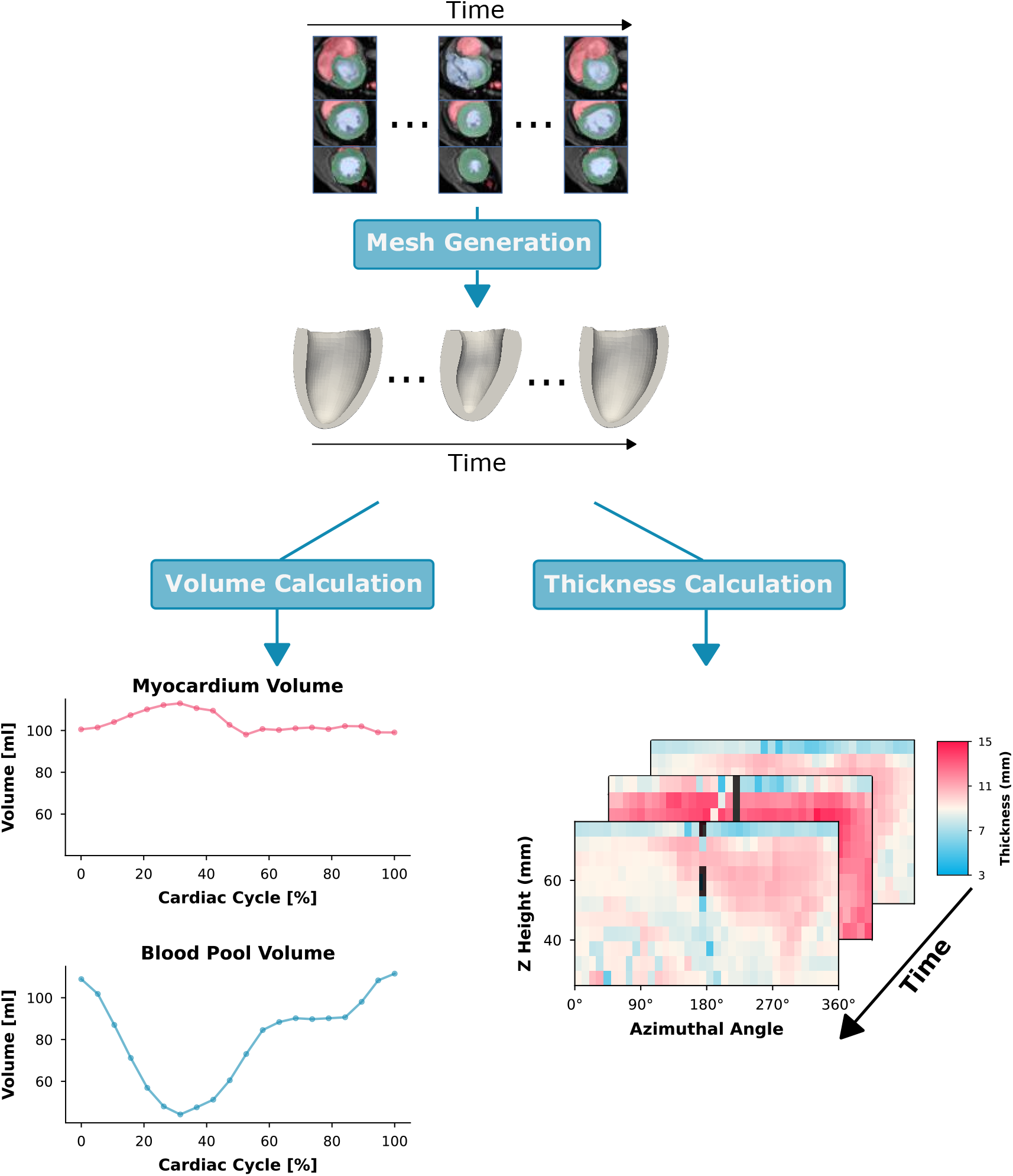
Workflow of the LVentiView Mesh Generation Module. **Top**: Segmented and cleaned short-axis MRI z-stacks acquired over the cardiac cycle (blue: blood pool, green: myocardium, red: right ventricle). **Middle**: Volumetric meshes fit to the MRI data across the cardiac cycle. **Bottom**: Quantitative outputs of the Mesh Generation Module, including blood pool and myocardial volumes as well as local myocardial wall thickness maps computed from the volumetric meshes throughout the cardiac cycle. In the thickness maps, black squares indicate invalid regions where the local wall thickness falls more than five standard deviations below the mean or where the value is undefined (NaN).

Myocardial volume is computed as the volume enclosed by the mesh using a discrete form of the divergence theorem, implemented in the vtkMassProperties.GetVolume() function from the Python library VTK. To calculate blood pool volume, the open end of the LV is capped to create a closed mesh, its total volume is computed, and the myocardial volume is subtracted.

Local myocardial wall thickness is derived from the fit mesh by first transforming the mesh node coordinates into cylindrical coordinates. The z-axis is defined by the longitudinal axis of the fitted mesh, which is aligned with the direction perpendicular to the SAX MRI slices. The nodes are then divided into 15 bins along the longitudinal axis. For bins, where the nodes form a ring-shaped geometry, the nodes are further subdivided azimuthally into 10° segments. This angular resolution was chosen to ensure a sufficient number of mesh nodes per bin for accurate thickness estimation, while maintaining sufficient spatial detail to resolve regional variations in wall thickness. Within each azimuthal bin, the radial myocardial thickness is calculated as the difference between the maximum and minimum radii. This approach produces a detailed map of local wall thickness along both the long axis and the circumferential direction of the LV.

While the workflow is fully automated by default, users can adjust mesh fitting parameters, select specific time steps, or enable/disable volume and thickness calculations.

### 3.4. GUI Software Implementation

LVentiView includes a GUI for automatic segmentation, 3D mesh-based LV reconstruction, and quantitative LV analysis described in Sections 3.1–3.3. We implemented the software in Python 3.12.3, with the GUI built using PyQt5 (v5.15.10 with Qt v5.15.2). The analysis backend relies on NumPy (v2.4.3), SciPy (v1.17.1), tqdm (v4.67.3), PyVista (v0.47.1), pydicom (v2.4.4), meshio (v5.3.5), imageio (v2.33.1), scikit-image (v0.26.0), pandas (v3.0.1), seaborn (v0.13.2), and PyTorch (v2.10.0+cu126). We converted the pretrained segmentation models from TensorFlow using onnx2torch (v1.5.15) into PyTorch.

We executed LVentiView on NVIDIA GeForce RTX 4060 GPU. Segmentation of a 4D MRI dataset with 20 time frames and 9 SAX slices per time frame required 12 s (≈ 0.07 s per slice). Fitting a mesh to one time step with 9 SAX slices took 2 min 44 s. Averaged across all 44 4D MRI data sets included in this study, the total mesh fitting time for the complete cardiac cycle was 58 min ± 9 min.

## 4. Experiments

### 4.1. Validation on Patient Data

We validate both modules of LVentiView on the publicly available Sunnybrook Cardiac Dataset (Radau, Lu, Connelly, Paul, Dick and Wright, 2009a,b), which comprises cine steady-state free precession MRI scans from 45 individuals covering the full cardiac cycle. MRI series were acquired in SAX view using a 1.5, T GE Signa MRI system during 10–15 s breath-holds. The dataset includes 12 patients with heart failure with infarction, 12 patients with heart failure without infarction, 12 with left ventricular hypertrophy, and 9 healthy controls, classified following the criteria of (Alfakih, Plein, Thiele, Jones, Ridgway and Sivananthan, 2003). One healthy subject (*SCD3901*) was excluded due to corrupted images, leaving 44 subjects for analysis. The data set includes manual annotations of SAX images at end-diastole (ED) for all 44 scans, provided by an experienced cardiologist and independently verified by a second expert.

#### 4.1.1. Automatic Segmentation

The Segmentation Module is evaluated by comparing LVentiView segmentation masks to the expert manual segmentations provided in the Sunnybrook Cardiac Dataset. Using inter-expert Dice scores of 0.93 for the blood pool and 0.88 for the myocardium as a reference (Bai et al., 2018), LVentiView achieves blood pool Dice scores at the inter-expert level and myocardium Dice scores slightly below (Fig. 3A). The average Dice scores for the blood pool were 0.93 ± 0.05 for healthy subjects, 0.95 ± 0.05 for patients with heart failure with infarction, 0.93 ± 0.07 for patients with heart failure without infarction, and 0.91 ± 0.06 for patients with left ventricular hypertension. For the myocardium, the corresponding Dice scores were 0.78 ± 0.09 for healthy subjects, 0.80 ± 0.10 for heart failure with infarction, 0.74 ± 0.14 for heart failure without infarction, and 0.77 ± 0.11 for left ventricular hypertension (Fig.3A).

**Figure 3:**
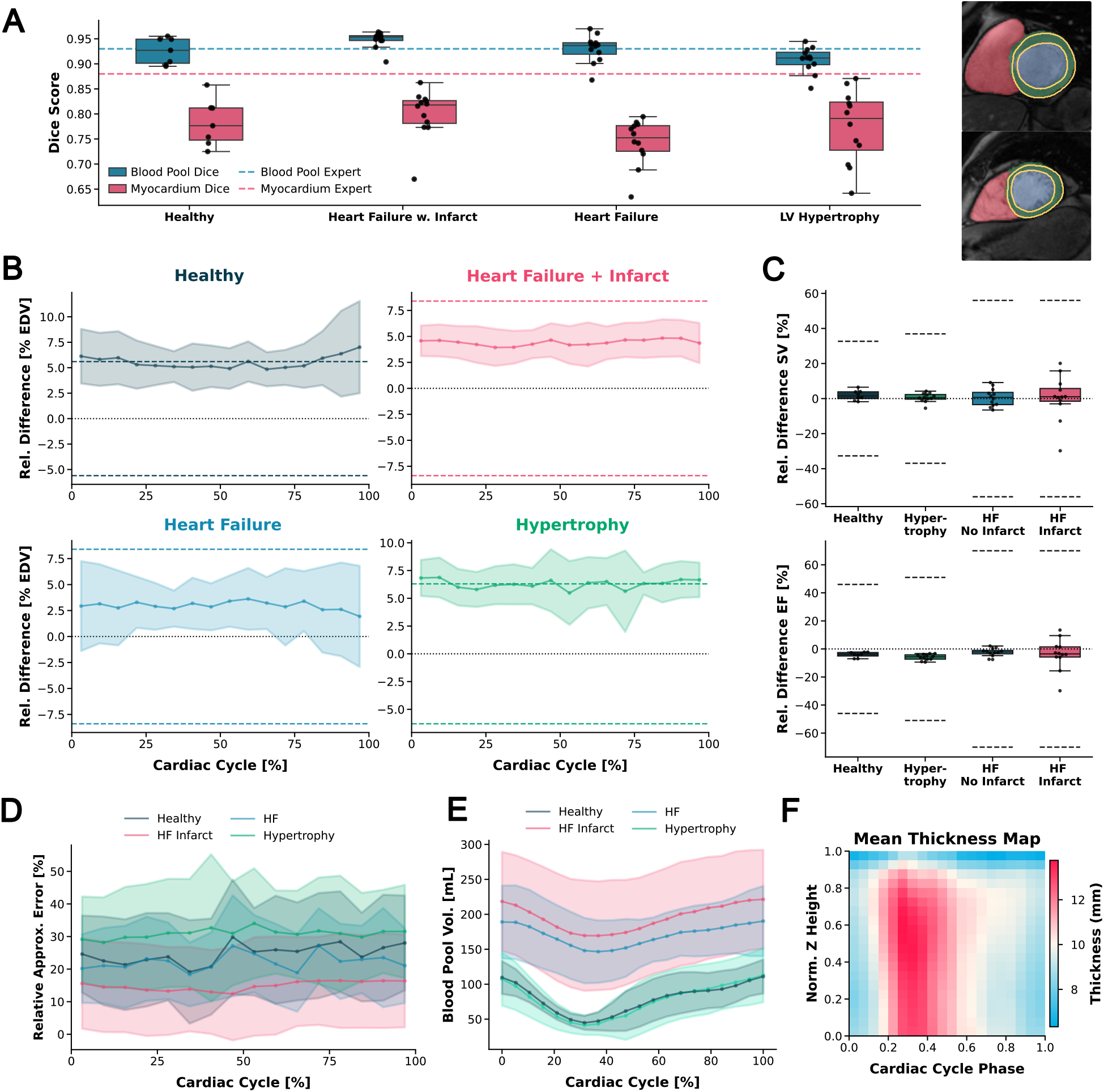
Verification of the LVentiView segmentation and mesh generation modules on patient data using SAX MRI only. **(A)** Left: Dice score distributions for blood pool and myocardium segmentations across four pathological groups. Right: Representative SAX MRI slices with manual (orange) and automated blood pool (blue) and myocardium (green) contours. Top: blood pool Dice= 0.97, myocardium Dice = 0.87. Bottom: blood pool Dice = 0.98, myocardium Dice = 0.85. **(B)** EDV-normalized relative difference between blood pool volumes computed from the sliced mesh and segmentation masks (*V*_*mesh*_ −*V*_*seg*_) for each pathology. In all line plots in this figure solid lines and shaded regions represent mean ± 1 SD across individual trajectories interpolated to 16 time steps. **(C)** Relative differences in SV (top) and EF (bottom) computed from the sliced mesh and segmentation masks. **(D)** Relative approximation error of Simpson’s method, defined as the relative difference between volumes computed directly from the mesh and from the sliced mesh. **(E)** Blood pool volume over the cardiac cycle for the four pathological groups. **(F)** Mean local myocardial thickness map averaged over eight healthy subjects. The x-axis denotes normalized cardiac cycle time and the y-axis LV wall thickness at each z-location averaged over all azimuthal angles. The z-axis corresponds to the LV longitudinal direction.

However, as manual annotations are used as the reference, these scores reflect agreement with human experts rather than absolute segmentation accuracy. Performance was consistent across all four pathologies, indicating that the Segmentation Module generalizes well without systematic bias toward any particular condition.

#### 4.1.2. Verification of Patient-Specific Meshes

To verify the Mesh Generation Module, blood pool volumes extracted from the sliced mesh via Simpson’s method are compared to those computed directly from the segmentation masks via Simpson’s method, with inter-expert variability used as the reference threshold. Given a reported inter-expert Dice variability of 7% (Bai et al., 2018) and assuming both experts are equally accurate, the expected relative volume difference between two expert segmentations is 14%.

Since absolute volume differences (Δ*V* ) remain approximately constant over the cardiac cycle while volumes decrease during systole, relative differences are normalized by the subject-specific end-diastolic volume (EDV) to avoid artificially inflated errors during systole (Fig. 3B, Supplementary Fig. S4A).

The inter-expert thresholds for this metric vary by pathology, as they depend on the EF. Assuming the worst case *V* = ESV, where the relative difference *ϵ* = Δ*V* /*V* is largest, the threshold for the EDV-normalized relative difference can be derived as Δ*V* /EDV = *ϵ*(1 − EF). Using pathology-specific ejection fractions following Alfakih et al. (2003) and substituting in the acceptable relative volume difference *ϵ* = 14%, the thresholds are 5.6% for healthy subjects, 8.4% for both types of heart failure, and 6.3% for hypertrophy. The mean EDV normalized relative differences were 5.6%±2.6% for healthy subjects, 2.9% ± 3.3% for heart failure without infarction, 4.4% ± 1.7% for heart failure with infarction, and 6.3% ± 2.0% for hypertrophy, all within or at the inter-expert threshold (Fig. 3B).

Stroke volume (SV) and EF derived from the sliced mesh showed differences well below the inter-expert threshold across all pathologies (Fig. 3C). The thresholds for SV and EF are derived assuming equal relative errors in EDV and ESV of *ϵ* = 14%, giving Δ*SV* /*SV* = *ϵ*(2 − EF)/EF and Δ*EF* /*EF* = (2*ϵ*)/EF.

In one case (*SCD3201*, hypertrophy), the initial mesh fitting produced a displaced and distorted mesh for a single time step (Supplementary Fig. S5A). Reinitializing the fitting from the mean of the successfully fitted meshes at neighboring time frames yielded a physiologically plausible result, with Dice scores improving substantially for both the myocardium (from 0.54 to 0.75) and the blood pool (from 0.36 to 0.75)(Supplementary Fig. S5B).

#### 4.1.3. Approximation Error of Simpson method

While Simpson’s method estimates LV volume by slicing the mesh and summing cross-sectional areas, the volume can alternatively be computed directly from the volumetric mesh using the VTK Python library. To confirm the accuracy of this direct computation, we validated it on an idealized LV geometry — a hemispherical shell with outer and inner diameters of 40 mm and 32 mm, meshed in COMSOL Multiphysics (v6.1.0.252) (COMSOL AB, 2024) with 2,630 nodes and 11,719 tetrahedral elements. The computed volume differed from the analytical ground truth by less than 1%, confirming negligible numerical error.

The volume computed directly from the mesh, however,differs from that obtained via Simpson’s method applied to the sliced mesh (Fig. 3D). We hypothesize that this discrepancy arises from known limitations of Simpson’s method: SAX MRI series do not fully cover the LV, and linear interpolation between slices does not capture the true patient-specific geometry (O’Dell, 2019). The mean relative differences between Simpson-derived and mesh-derived volumes were 25% ± 14% for healthy subjects, 23% ± 11% for heart failure without infarction, 15% ± 14% for heart failure with infarction, and 30% ± 14% for hypertrophy (Fig. 3D, Supplementary Fig. S4B–C). These differences are larger than the inter-expert threshold, indicating that volume calculation using Simpson’s method introduces inaccuracies.

#### 4.1.4. Physiological Plausibility of Generated Meshes

Having established that the meshes agree with segmentation-derived volumes within inter-expert variability, we examine their geometry and dynamics in more detail. For all four pathologies, volumes computed directly from the volumetric meshes increased during the ED state and decreased during the ES state, consistent with expected cardiac physiology (Bonow, Mann, Zipes and Libby, 2011). Myocardial volume showed only minor variation throughout the cardiac cycle, whereas blood pool volume changed substantially (Fig. 3B, Supplementary Fig. S6A), in agreement with recent studies (Kumar, Ryu, Manduca, Rao, Gibbons, Gersh, Chandrasekaran, Asirvatham, Araoz, Oh et al., 2021). Additionally, LVV was elevated throughout the cardiac cycle in heart failure pathologies compared with left ventricular hypertrophy and healthy subjects, consistent with previous reports of increased EDV and ESV in heart failure (Stoll, Hess, Rodgers, Bissell, Dyverfeldt, Ebbers, Myerson, Carlhaell and Neubauer, 2019).

#### 4.1.5. Regional Myocardial Thickness

The meshes additionally captured physiologically meaningful variations in regional myocardial wall thickness throughout the cardiac cycle for all four pathologies. As expected, the myocardial wall thickened during systole and thinned during diastole (Bonow et al., 2011, Fig. 3C, Supplementary Fig. S6B-–E). Subjects with left ventricular hypertrophy exhibited increased myocardial wall thickness (11.87 mm ± 1.35 mm) compared to the remaining groups (10.18 mm ± 1.14 mm), consistent with its established role as a diagnostic marker for hypertrophy (Baltabaeva et al., 2008, Fig. S6E). This demonstrates that the volumetric meshes created with LVentiView capture pathology-specific regional wall thickness that is not accessible from global volumetric indices alone.

## 5. Discussion

Patient-specific cardiac modeling requires accurate threedimensional representations of the left ventricle, yet existing segmentation platforms such as 3D Slicer and ITK-SNAP (Fedorov et al., 2012; Yushkevich et al., 2006) lack mesh generation capabilities, while cardiac simulation frameworks such as lifex (Africa, 2022) do not provide an image-to-mesh pipeline. LVentiView addresses this gap by providing a fully automated pipeline from MRI segmentation to patient-specific 3D LV mesh reconstruction, enabling regional myocardial thickness analysis across the full cardiac cycle. In the following, we discuss the performance of each pipeline component, the physiological plausibility of the generated meshes, and the volumetric approximation error inherent to Simpson’s method.

### 5.1. Automated Segmentation

For automatic segmentation to be clinically useful, it must perform at least as well as an expert human annotator. The inter-expert Dice score therefore represents a natural benchmark: if the variability between the automatic segmentation and a manual annotation matches the variability between two independent experts, the tool is performing at the level of a human annotator.

The Segmentation Module achieved blood pool Dice scores at the inter-expert level and myocardial Dice scores slightly below inter-expert level. The performance of the model is limited by the manual annotations it was trained on,which inherently reflect inter-expert variability in MRI segmentations. Consequently, reaching inter-expert-level blood pool Dice scores represents the best performance the model can achieve, and the slightly lower myocardium scores fall within the expected range. It should be noted that these scores reflect agreement with human expert annotations rather than absolute segmentation accuracy. Determining the true segmentation accuracy would require realistic synthetic MRI images with known ground-truth geometries, against which the automated segmentation could be objectively evaluated.

Nevertheless, by producing reproducible segmentation masks consistently across all four pathologies, LVentiView provides a reliable foundation for downstream mesh generation and volumetric analysis.

### 5.2. Verification of Patient-Specific Meshes

The patient-specific meshes were considered verified if the volume difference between the sliced mesh and the segmentation masks fell within the range of inter-expert variability. This threshold was derived from the expected volume difference between two independent experts segmenting the same image using Simpson’s method.

The average agreement between the sliced mesh and the segmentation masks falls within one standard deviation of the inter-expert Dice scores. This geometric agreement translates directly to the volumetric metrics: the EDV-normalized relative volume differences, as well as the derived SV and EF differences, all fell within the thresholds derived from inter-expert variability across all four pathologies. This confirms that volumes and cardiac parameters computed from the patient-specific meshes are as accurate as those obtained from expert manual segmentations.

### 5.3. Robustness of Mesh Fitting

Mesh fitting from an initial average LV geometry succeeded for all subjects and time steps with one exception. In one hypertrophic subject (*SCD3201*), mesh fitting failed for a single systolic time step, producing a distorted and displaced mesh. We attribute this to a missing MRI slice at the LV base, which led to insufficient information for the fitting to converge to a physiologically plausible solution. Incomplete basal slice coverage is not uncommon in clinical MRI acquisitions.

Reinitializing the fit from the mean of the successfully fitted meshes at the neighboring time points resolved the issue, yielding a geometrically plausible mesh with substantially improved Dice scores for both the myocardium (from 0.54 to 0.75) and the blood pool (from 0.36 to 0.75). This suggests that when mesh fitting produces unrealistic results for a single time step, initialization from neighboring time frames is a robust fallback strategy.

### 5.4. Approximation Error of Simpson method

We hypothesize that the large difference between volumes calculated directly from the mesh and from the sliced mesh via Simpson’s method highlights the approximation error inherent to Simpson’s method. Simpson’s method systematically underestimates the volume calculated directly from the mesh. Since the direct volume calculation was validated against an idealized geometry with a known analytical ground truth, we consider the directly computed mesh volume to be an accurate representation of the true LV volume.

This discrepancy therefore could point to approximation errors in Simpson’s method, consistent with findings reported by O’Dell (2019), which identify incomplete basal and apical slice coverage in SAX MRI series and the assumption of linear geometry between slices as likely sources of this error.

This finding has an important clinical implication: volumes derived from Simpson’s method, which is the clinical standard, underestimate the true LV volume. We found relative differences of 25.08 ± 13.96% for healthy subjects,22.64 ± 11.39% for heart failure without infarction, 14.93 ± 13.95% for heart failure with infarction, and 30.74 ± 14.40% for hypertrophy. This error is larger than the error introduced by variability between expert annotators. The volumetric meshes generated by LVentiView and direct volume computation thereof could therefore improve the accuracy of LV volume estimation beyond what is currently achievable with Simpson’s method in clinical practice.

### 5.5. Limitations

Several limitations of LVentiView should be acknowledged. First, the accuracy of the Segmentation Module was assessed by comparison against expert manual annotations, which quantifies agreement with human experts rather than true segmentation accuracy. As discussed above, determining true accuracy would require realistic synthetic MRI images with known ground-truth geometries.

Second, LVentiView currently operates on SAX MRI slices only. Relying exclusively on SAX slices introduces ambiguity in the precise location of the basal and apical planes, which may lead to inaccurate mesh shapes in regions beyond the coverage of the SAX stack. Incorporating longaxis (LAX) MRI slices into the mesh fitting procedure could mitigate this limitation by providing complementary information about the LV extent.

Third, mesh fitting requires approximately 3 minutes per time frame, amounting to an average of 58 minutes to fit a full 4D MRI dataset on an NVIDIA GeForce RTX 4060 GPU. While this is feasible for research use, computation time scales with the number of time frames and depends on the available hardware, which may limit throughput in clinical settings.

Finally, LVentiView is currently optimized for cardiac MRI and does not yet support cardiac computed tomography (CT). Extending support to CT would require an additional segmentation model capable of segmenting the LV from CT images, after which the existing mesh fitting procedure could be applied.

LVentiView is openly available and we encourage users to contribute to its continued development.

## 6. Conclusion

Three-dimensional reconstruction of the left ventricle from cardiac MRI supports both clinical diagnosis and patient-specific cardiac simulations. In this work, we presented LVentiView, an open-source software package that unifies automated MRI segmentation, 3D mesh-based LV reconstruction, volumetric analysis, and regional myocardial thickness calculation in a single reproducible workflow. Validated on a publicly available cardiac MRI dataset, LVentiView achieved blood pool segmentation performance at the inter-expert level. Volumes derived from the patientspecific meshes were shown to be as accurate as those obtained from expert manual segmentations, confirming the reliability of the meshes generated by the pipeline. With its accessible graphical interface requiring no expert knowledge and open-source availability on GitHub https://github.com/InaBraun01/LVentiView.git, LVentiView lowers the barrier between clinical MRI data and patient-specific biomechanical modeling, while enabling more accurate cardiac parameter estimation than is achievable with Simpson’s method alone.

## Supporting information

Full Supplementary Materials

## Declaration of generative AI use

During the preparation of this work the authors used Claude (Anthropic) and ChatGPT (OpenAI) in order to assist with GUI implementation, code cleaning, plotting, as well as for manuscript editing and rephrasing. After using these tools, the authors reviewed and edited the content as needed and take full responsibility for the content of the published article.

## Acknowledgements

The authors would like to thank Constanze Schmidt for her thoughtful comments and suggestions that helped improve the manuscript. This research was conducted within the Max Planck School Matter to Life, supported by the Dieter Schwarz Foundation in collaboration with the Max Planck Society. This research was supported by the DZHK (German Centre for Cardiovascular Research), funding code: 81X3300402.

## CRediT authorship contribution statement

**Ina Braun:** Software, Data curation, Formal analysis, Writing – original draft. **Yong Wang:** Conceptualization, Writing - review editing. **Alexander S. Ecker:** Conceptualization, Writing - review editing. **Eberhard Bodenschatz:** Supervision, Conceptualization, Funding acquisition, Writing - review editing.

## Footnotes

1 https://github.com/InaBraun01/LVentiView.git

