## Supplementary material for "LVentiView: An Open-Source Software for Automated 3D Left Ventricular Mesh Reconstruction and Analysis from Cardiac MRI": Full Supplementary Materials

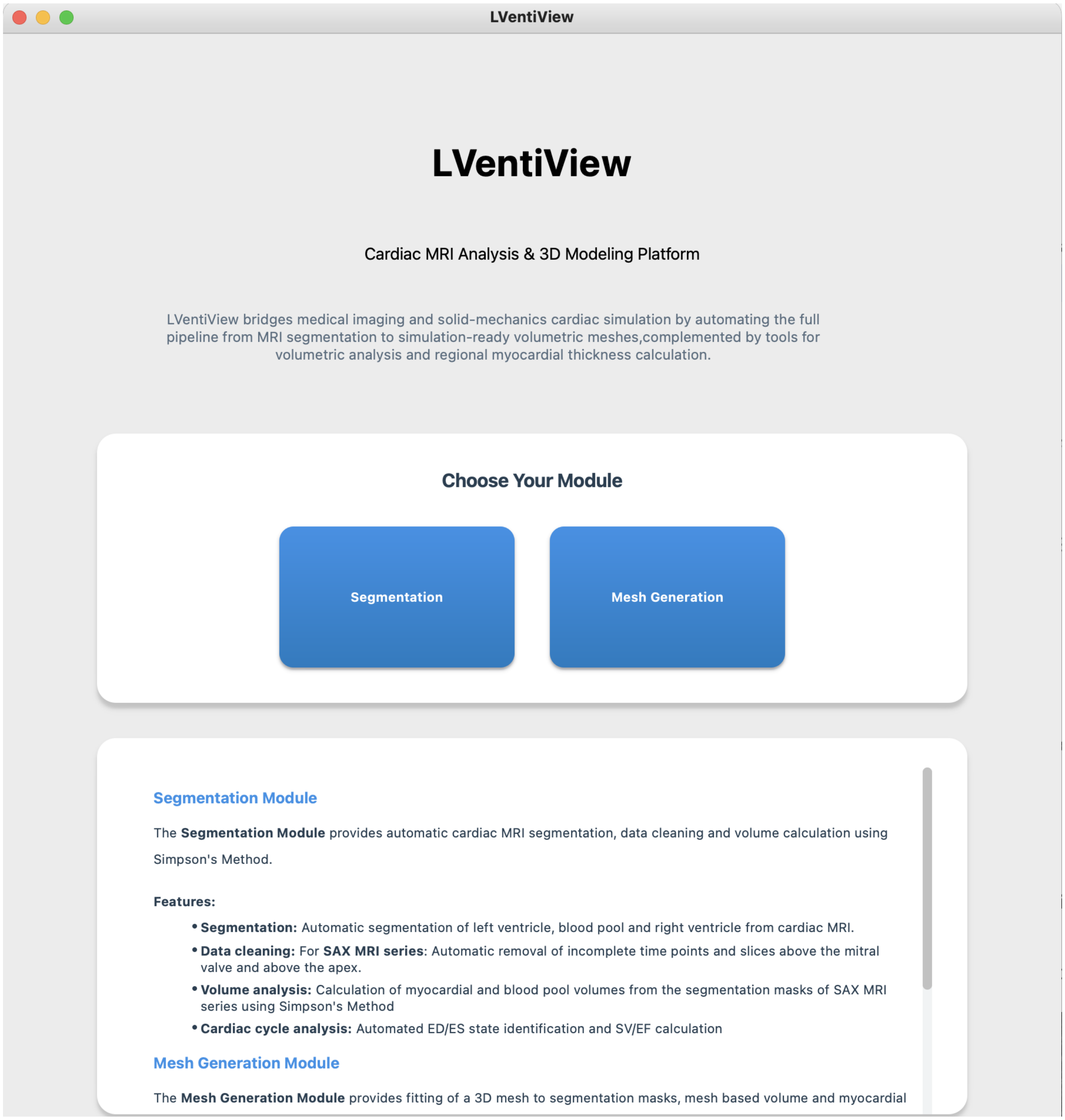


Figure S1: Start Page of LVentiView. Overview of the LVentiView software interface. The blue Segmentation and Mesh Generation buttons provide access to the corresponding modules. A brief summary of the available modules is displayed below the navigation buttons.

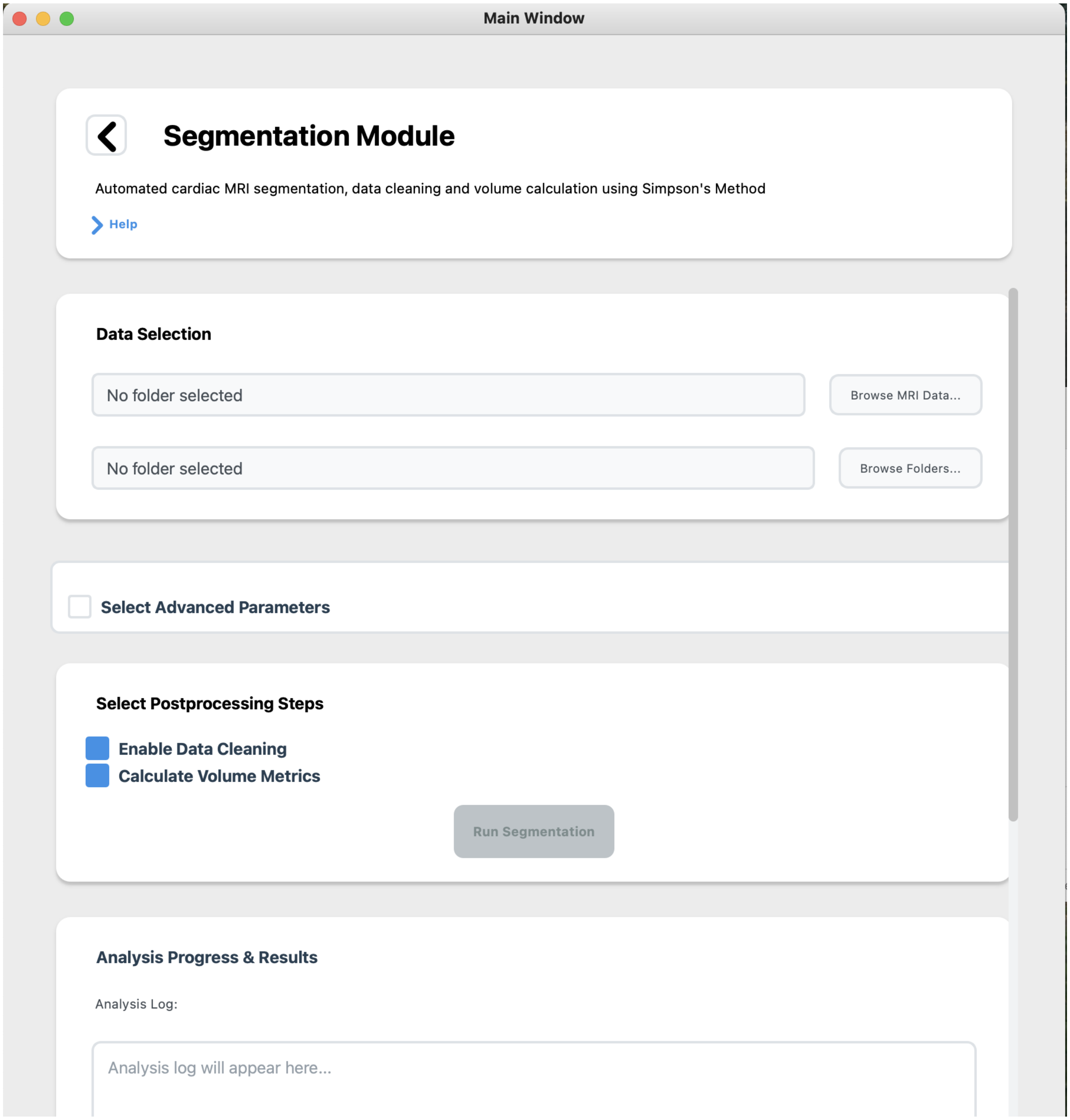


Figure S2: Segmentation Module of LVentiView. User interface of the Segmentation Module. The help section provides detailed information on the workflow and parameters. Users can select the MRI series to be segmented and the output directory, optionally enable advanced parameters, and choose post-processing steps such as data cleaning and volume computation using Simpson's method. Once the required inputs are selected, segmentation can be started, and the resulting outputs are displayed below in the interface.

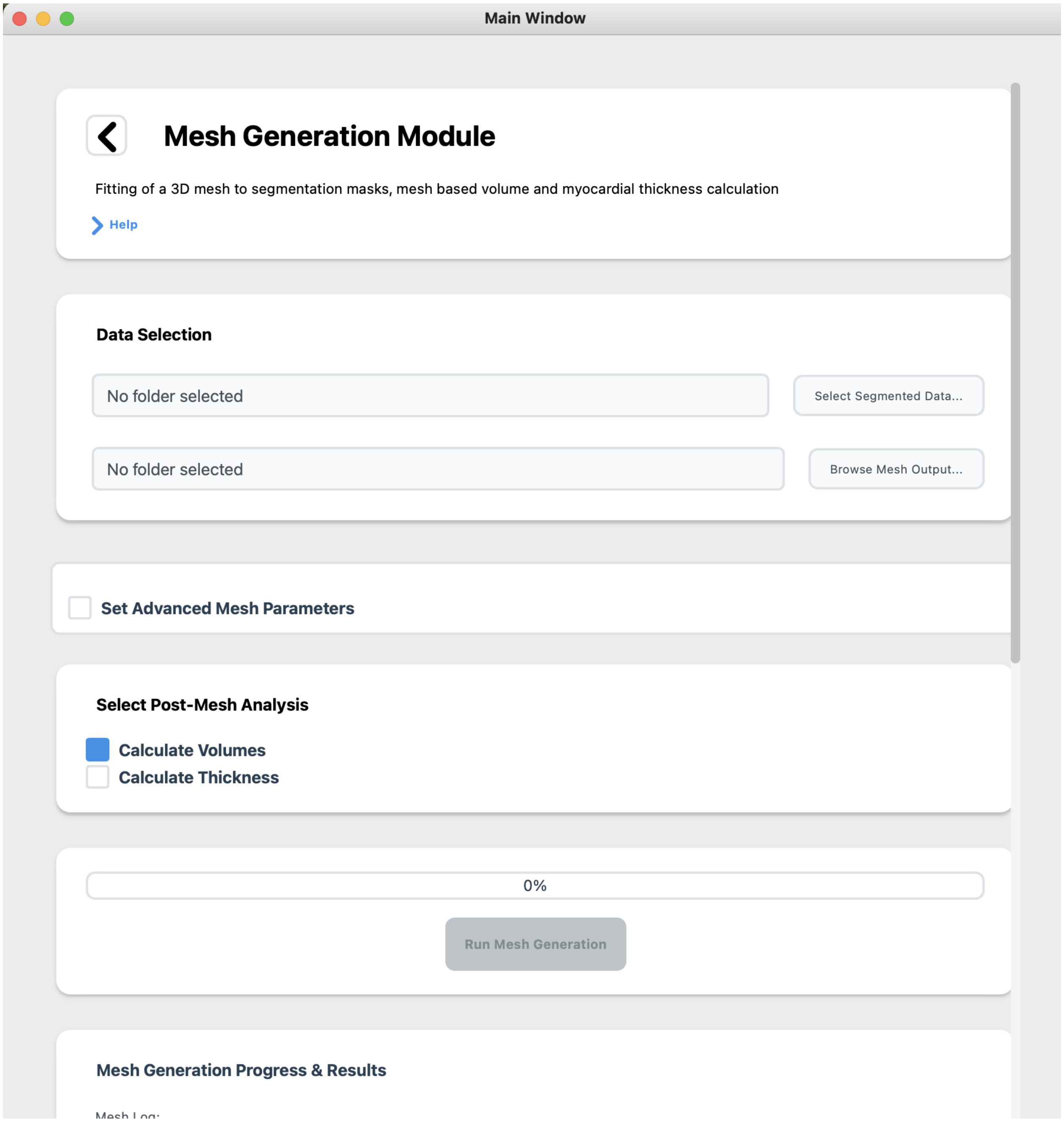


Figure S3: Mesh Generation Module of LVentiView. User interface of the Mesh Generation Module. The help section describes the fitting procedure and available parameters. Segmented MRI data and an output directory can be selected, advanced parameters can be enabled, and post-processing options such as volume and local thickness computation can be chosen. Mesh fitting progress is shown via a progress bar, with the resulting volumetric meshes and derived quantities displayed below.

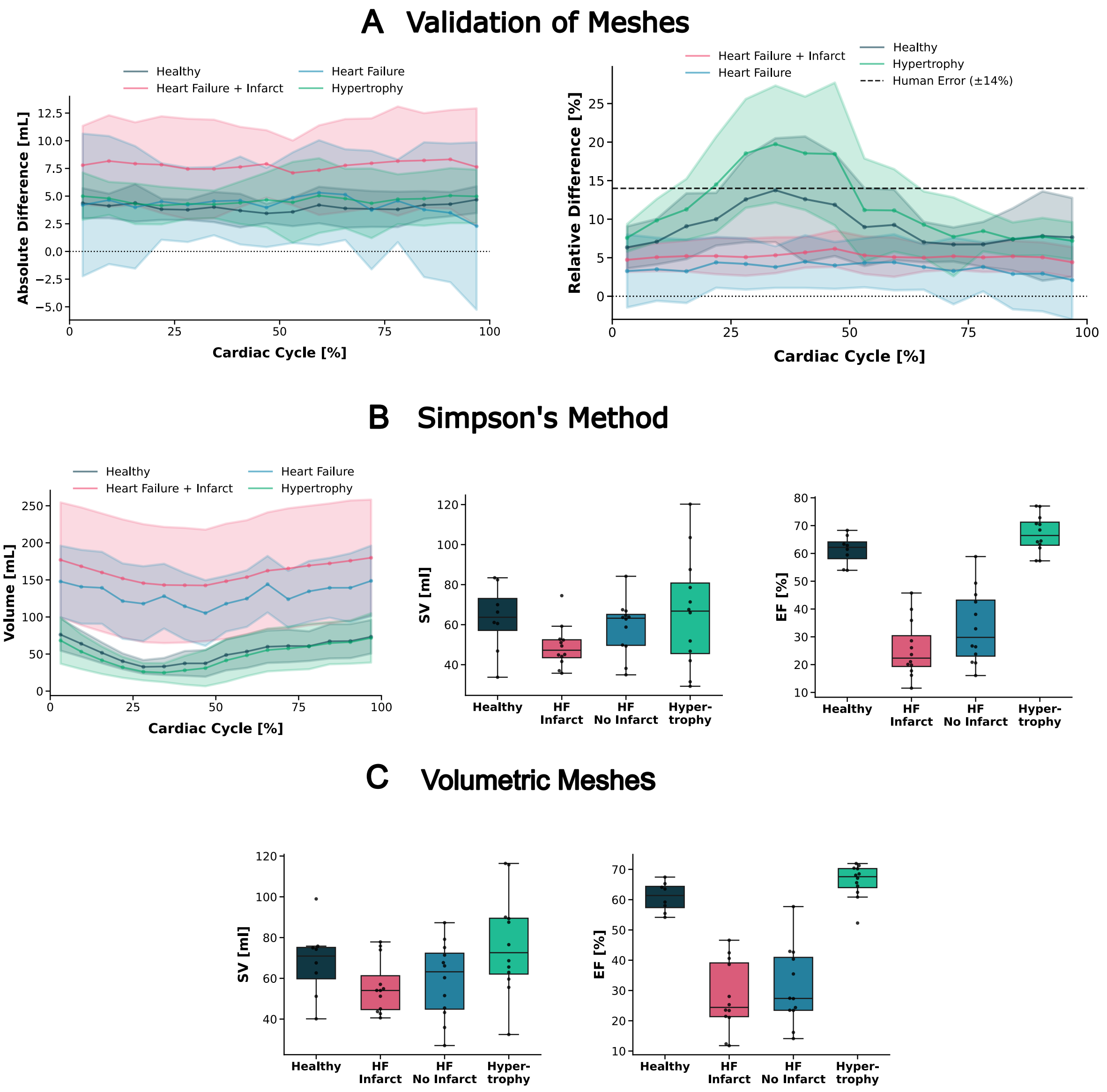


Figure S4: Verification of the LVentiView Segmentation and Mesh Generation Modules on patient data using SAX MRI only. **(A)** Difference between blood pool volumes computed via Simpson's method from the sliced mesh and from the segmentation masks ($V_{mesh}$ - $V_{seg}$), shown as absolute (left) and relative (right) differences over the cardiac cycle. **(B)** Cardiac parameters computed via Simpson's method from the segmentation masks for the four pathological groups: blood pool volume over the cardiac cycle (left), stroke volume (middle), and ejection fraction (right). **(C)** Corresponding parameters computed directly from the volumetric meshes fit to SAX MRI series.

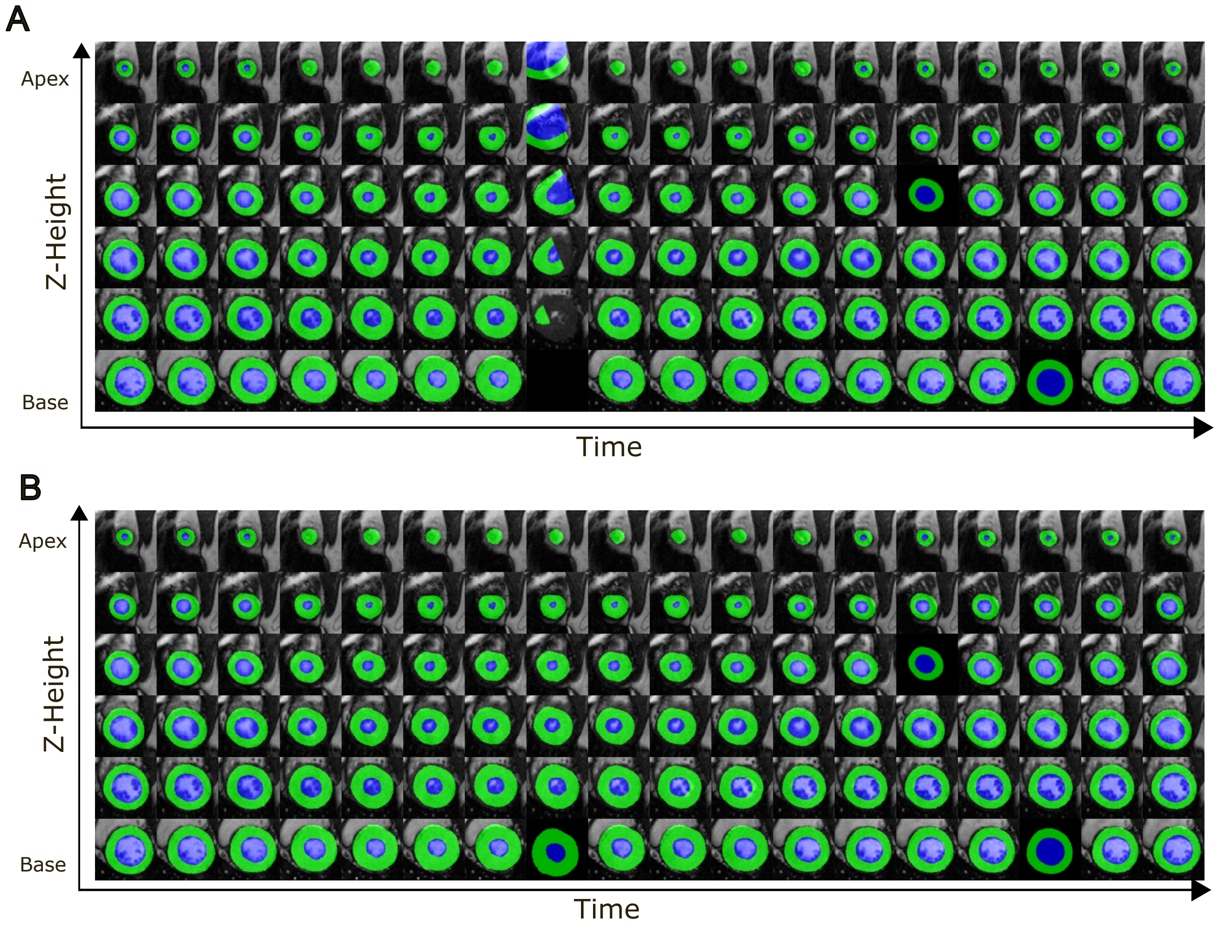


Figure S5: Mesh fitting failure and recovery for subject *SCD3201* (hypertrophy). Sliced meshes overlaid on SAX MRI images across the full cardiac cycle, with the mesh wall shown in green and the inside of the mesh shown in blue. **(A)** Initial mesh fitting, where one time step produced a displaced and distorted mesh. **(B)** Corrected result after reinitializing the fitting for the affected time step from the mean of the successfully fitted meshes at neighboring time frames.

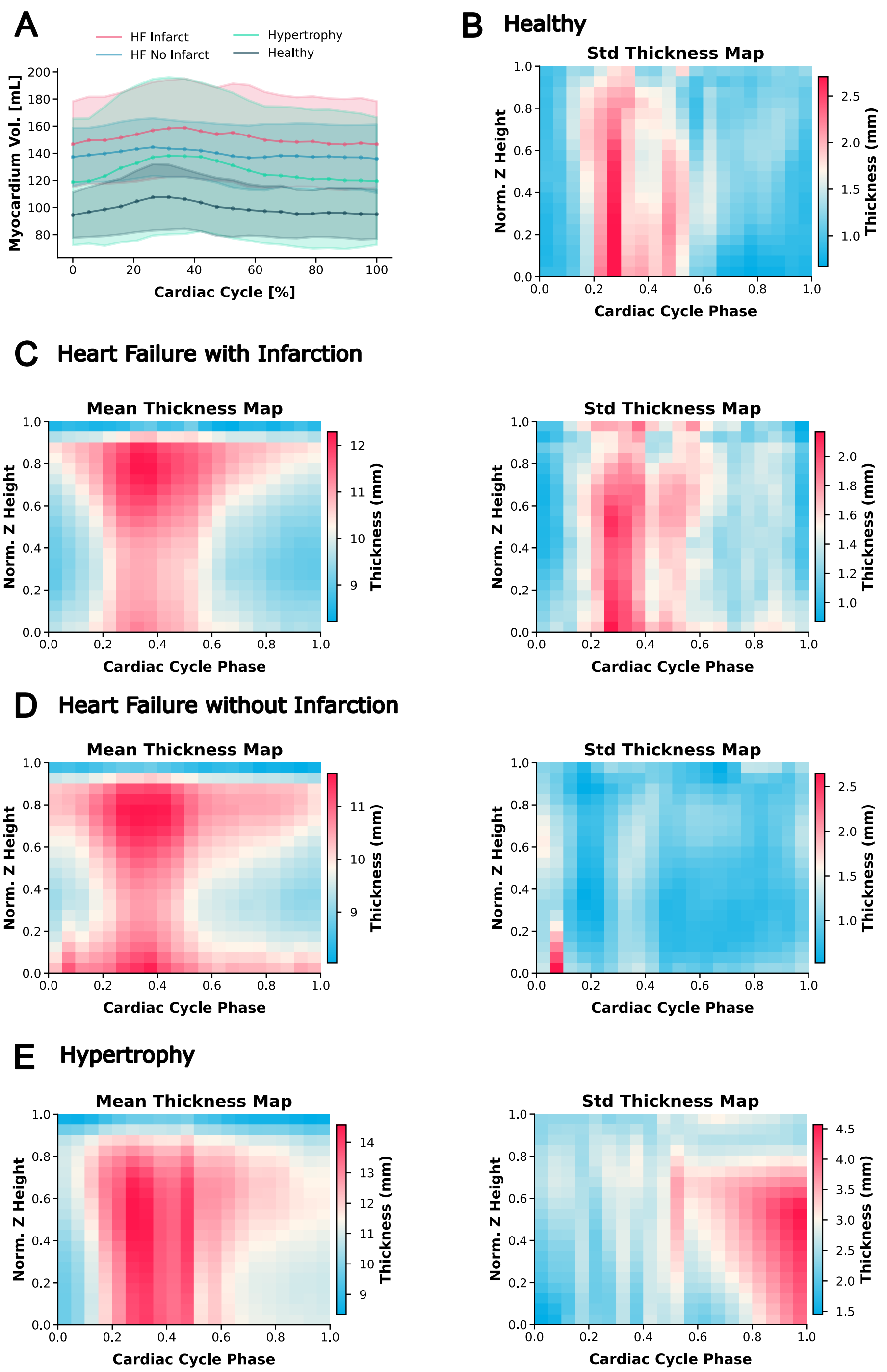


Figure S6: Physiological consistency of volumetric meshes generated from SAX MRI. **(A)** Myocardial volume trajectories over the cardiac cycle for the four pathological groups. The mean myocardial volume is shown as a solid line, and shaded regions indicate one standard deviation. To compute the average myocardial volume for each pathology, individual trajectories were interpolated to 20 time steps over the cardiac cycle and averaged pointwise. **(B)** Standard deviation of local myocardial wall thickness for healthy anatomies. **(C–E)** Mean and standard deviation of local myocardial wall thickness for patients with heart failure with infarction **(C)**, heart failure without infarction **(D)**, and left ventricular hypertrophy **(E)**. In all thickness maps, the x-axis represents normalized cardiac cycle time and the y-axis represents wall thickness at a given z-location, averaged over all azimuthal angles.
